## Supplementary information for "Informative and adaptive distances and summary statistics in sequential approximate Bayesian computation"

### 1 Optimal summary statistics to recover distribution features

**Theorem 1.** *Denote the joint distribution of parameters and data* $\Theta, Y \sim \pi(\theta, y)$, *with prior marginal* $\pi(\theta) = \int \pi(\theta, y)\, dy$, *likelihood* $\pi(y|\theta) = \pi(\theta, y)/\pi(\theta)$, *and posterior* $\pi(\theta|y) = \pi(\theta, y)/\pi(y) = \pi(y|\theta)\pi(\theta)/\pi(y)$. *Given a parameter transformation* $\lambda : \mathbb{R}^{n_\theta} \to \mathbb{R}^{n_\lambda}$ *such that* $\mathbb{E}_{\pi(\theta)}[|\lambda(\theta)|] < \infty$, *define summary statistics as the conditional expectation*

$$s(y) := \mathbb{E}[\lambda(\Theta)|Y = y] = \int \lambda(\theta)\pi(\theta|y)\, d\theta\,.$$

*Given observed data* $y_{obs}$, *acceptance threshold* $\varepsilon$, *and assuming the distance metric* $d(s(y), s(y_{obs})) = \|s(y) - s(y_{obs})\|$ *is norm-induced, denote the ABC posterior distribution*

$$\pi_{\text{ABC},\varepsilon}(\theta|s(y_{obs})) \propto \int I[\|s(y) - s(y_{obs})\| \leq \varepsilon]\pi(y|\theta)\, dy \cdot \pi(\theta).$$

*Then, it holds*

$$\left\|\mathbb{E}_{\pi_{\text{ABC},\varepsilon}}[\lambda(\Theta)|s(y_{obs})] - s(y_{obs})\right\| \leq \varepsilon, \tag{1}$$

*and therefore*

$$\lim_{\varepsilon \to 0} \mathbb{E}_{\pi_{\text{ABC},\varepsilon}}[\lambda(\Theta)|s(y_{obs})] = \mathbb{E}[\lambda(\Theta)|Y = y_{obs}]. \tag{2}$$

*Proof.* Based on Fearnhead and Prangle [2012] and Jiang et al. [2017], a simple extension of the argumentation in the latter. Note that $s(y)$ is almost surely finite due to $\mathbb{E}[|\lambda(\theta)|] < \infty$ and Fubini's Theorem. As for the induced $\sigma$-algebras holds $\sigma(s(Y)) \subset \sigma(Y)$, $s(Y)$ is also a version of the conditional expectation $\mathbb{E}[\lambda(\Theta)|s(Y)]$, since

$$s(Y) = \mathbb{E}[s(Y)|s(Y)] = \mathbb{E}[\mathbb{E}[\lambda(\Theta)|Y]|s(Y)] = \mathbb{E}[\lambda(\Theta)|s(Y)]$$

by, respectively, measurability, definition, and tower property. Thus, denoting the acceptance region

$$A = \{\|s(Y) - s(y_{\text{obs}})\| \leq \varepsilon\} \in \sigma(s(Y)),$$

with $\mathbb{E}[\lambda(\Theta)|A] = \mathbb{E}[\lambda(\Theta)\mathbb{1}_A]/\mathbb{E}[\mathbb{1}_A]$, we have

$$\mathbb{E}_{\text{ABC},\varepsilon}[\lambda(\Theta)|s(y_{\text{obs}})] = \mathbb{E}[\lambda(\Theta)|A] = \mathbb{E}[s(Y)|A],$$

such that by Jensen's inequality, given convexity of the norm,

$$\left\|\mathbb{E}_{\pi_{\text{ABC},\varepsilon}}[\lambda(\Theta)|y_{\text{obs}}] - s(y_{\text{obs}})\right\| = \|\mathbb{E}[s(Y) - s(y_{\text{obs}})|A]\| \leq \mathbb{E}[\|s(Y) - s(y_{\text{obs}})\| \, |A] \leq \varepsilon.$$

(2) then follows directly from (1) by definition of $s(y_{\text{obs}})$. □

Therefore, e.g. for $\lambda(\theta) = (\theta^1, \ldots, \theta^k)$, the corresponding first $k$ moments of the true posterior distribution are recovered by an ABC analysis employing the posterior expectation $s(y) = \mathbb{E}[\lambda(\Theta)|Y = y]$ as summary statistic, for $\varepsilon \to 0$. For $k \to \infty$, $\varepsilon \to 0$, and e.g. assuming existence of moment-generating functions, the approximate posterior converges to the true posterior.

### 2 Effective sample sizes

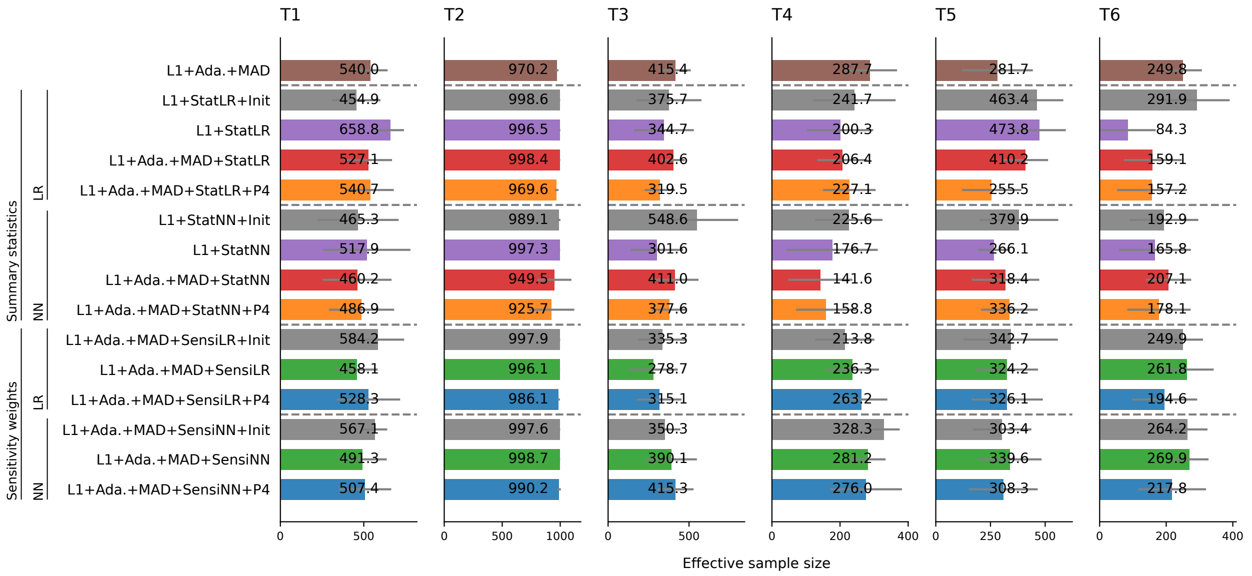


Figure S1: Effective sample sizes (ESS) for models T1-6. Given particles $P_{n_t} = \{(\theta_i, w_i)\}_i$ accepted in the last generation, the ESS is defined as $\text{ESS} = (\sum_i w_i)^2 / \sum_i w_i^2$ [Martino et al., 2017]. Shown are means and standard deviations (grey error bars) over all performed runs.

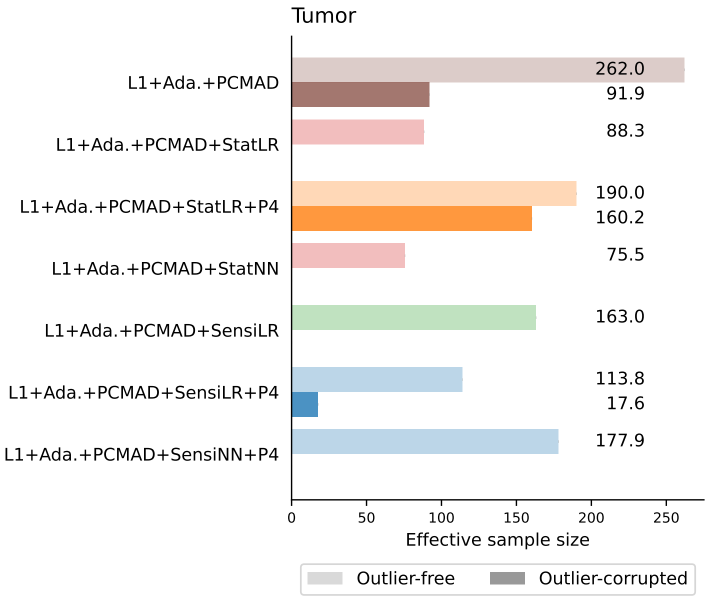


Figure S2: Effective sample sizes (ESS) for the tumor model, on outlier-free (light bars) and outlier-corrupted (dark bars) data, for selected settings. Note that on outlier-corrupted data, only three settings were run.
